# Neural Dynamics of Automatic Speech Production

**DOI:** 10.64898/2026.02.10.705088

**Authors:** Amirhossein Khalilian-Gourtani, Chenqian Le, Faxin Zhou, Erika Jenson, Patricia Dugan, Orrin Devinsky, Werner K. Doyle, Daniel Friedman, Yao Wang, Adeen Flinker

**Affiliations:** Neurology Department, NYU Grossman School of Medicine; Electrical Engineering Department, New York University; Biomedical Engineering Department, New York University; Neurosurgery Department, NYU Grossman School of Medicine

## Abstract

Speech is a defining human behavior, and this ability depends critically on speech motor cortex. While the ventral precentral and postcentral gyri are classically regarded as chiefly articulatory and somatosensory regions, a growing body of literature challenges this simplification. Most prior research, however, has examined cued or structured speech production tasks, neglecting the automatic, overlearned speech commonly utilized in clinical assessment. Consequently, the neural dynamics and precise timing of cortical recruitment during automatic speech remain poorly understood. Here, we present intracranial electrocorticography (ECoG) recordings from the left perisylvian cortex in participants performing automatic speech such as counting and recitation of overlearned sequences. We investigate neural dynamics using encoding (multivariate temporal response function) and decoding (deep neural network speech synthesis) models. We show that automatic speech engages a distributed network across superior temporal, precentral, and post-central cortices, characterized by attenuated pre-articulatory activity and weaker frontal encoding. Furthermore, two complementary decoding strategies reveal that speech motor cortex represents a mixture of feedforward and feedback signals, with a subset of sites exhibiting exclusively feed-forward dynamics. These results delineate the spatiotemporal cortical organization of automatic speech and establish that the speech motor cortex supports more complex dynamics than purely feedforward control.

## Introduction

Speech production is an extraordinarily rapid and demanding motor behavior, requiring accurately timed coordination across more than a hundred laryngeal, orofacial, and respiratory muscles whose cortical representations lie within ventral sensorimotor cortex (vSMC) (***Simonyan and Horwitz (2011)***; ***Guenther (2016)***). Much of how we think about this brain region still traces back to Penfield’s classical homunculus, which drew a sharp boundary between motor and sensory cortices along the pre- and post-central gyri and proposed a clean, orderly somatotopy linking cortical locations to adjacent body parts (***Penfield and Boldrey (1937)***; ***Catani (2017)***). While this framework has been foundational, a growing literature in human speech production is steadily revising this picture. Studies using fMRI, and ECoG recordings now show that the functional organization of vSMC during speech is more distributed, overlapping, and context-dependent than the traditional model suggests (***Hsu et al. (2025)***; ***Conant et al. (2018)***; ***Bouchard et al. (2013)***; ***Cheung et al. (2016)***). Speech-related activity spans precentral and postcentral regions and shows substantial co-activation before and during speech production, consistent with dynamic interactions between motor planning, execution, and sensory feedback (***Grabski et al. (2012)***; ***Eichert et al. (2020)***). Collectively, these findings challenge a strictly modular homuncular view and instead point to a dynamic and task-dependent organization of vSMC in speech (***Simonyan et al. (2016)***; ***Conant et al. (2014)***).

Most human speech studies have emphasized task-based or cued speech production, such as reading aloud, picture naming, word generation, or sentence repetition (***Towle et al. (2008)***; ***Haller et al. (2018)***; ***Saravani et al. (2019)***; ***Khalilian-Gourtani et al. (2024)***; ***Woolnough and Tandon (2024)***; ***Esmaeili et al. (2026)***). These tasks reliably engage the distributed language network and have been heavily used in speech decoding and brain-computer interface applications (***Chen et al. (2024)***; ***Wilson et al. (2020)***; ***Chakrabarti et al. (2015)***; ***Luo et al. (2022)***; ***Duraivel et al. (2023)***; ***Wyse-Sookoo et al. (2024)***; ***Metzger et al. (2023)***). In contrast, automatic speech—defined as the production of overlearned, highly practiced sequences like counting, reciting months, or repeating familiar prose—appears to engage this network differently. In a PET study, automatic tasks that simply repeated phoneme sequences or rote lists did not reliably activate classic language cortices, and only the recitation of memorized prose produced clear left-lateralized increases in articulation and auditory processing regions (***Bookheimer et al. (2000)***). This suggests that automatic speech may have a reduced demand on lexical retrieval and controlled planning relative to cued language tasks (***Gan et al. (2013)***). Indeed, imaging studies comparing automatic sequences (e.g., counting) with more controlled verbal fluency show differential hemispheric and regional involvement, with controlled tasks preferentially engaging frontal and temporal language regions (***Birn et al. (2010)***).

Clinically, the distinction between automatic and cued task-based speech has important ramifications for stimulation mapping of eloquent cortex. During awake neurosurgery, direct cortical stimulation is used to identify sites critical for speech and language by transiently disrupting performance, and the choice of behavioral task can influence the type of impairment observed (***de Camargo et al. (2025)***; ***Arya et al. (2018)***; ***Michalak et al. (2025)***). Classical mapping protocols have often relied on automatic speech tasks, such as counting or reciting days of the week, to establish stimulation induced impairments (i.e. motor and speech arrest) because these behaviors are easy to elicit, intereprt, and robust under stimulation. More demanding cued based language tasks, including word reading or picture naming, are followed when automatic speech fails to reveal a stimulation effect (***Michalak et al. (2025)***). Despite their widespread clinical use, the neural dynamics and spatiotemporal recruitment of perisylvian cortex during automatic speech remain poorly characterized, particularly using high-resolution methods such as electrocorticography (ECoG), which are well suited to resolve rapid speech-related cortical activity.

Here, we used electrocorticography recordings from neurosurgical patients to investigate the cortical recruitment and neural dynamics during automatic speech production. Using a temporal encoding model of neural activity, we quantified the cortical regions that are engaged during automatic speech tasks. We investigated the temporal dynamics of recruitment across early preparatory activations in motor, precentral, and frontal regions, as well as how the timing of encoding is organized across perisylvian cortex. Further, using a novel deep neural network architecture we showed how automatic speech can significantly be decoded from neural activity. The decoding performance critically depended on electrodes localized to speech-motor and auditory areas. By constraining the timing causality of our decoding architecture, we delineated the mixture of feedforward and feedback signals in speech motor cortex. This characterization of automatic speech dynamics offers new insights into functional mapping for neurosurgical planning and refining models of the speech production system.

## Results

We collected intracranial recordings sampling left perisylvian cortex in participants (*N* = 7) producing automatic and prompted cue-based speech tasks (see Participant information). Participants counted aloud from one to twenty at their own pace without pauses (counting task) as well as recited the days of the week and the months of the year (days + months task). These automatic speech tasks (Figure 1A) did not require any visual or auditory cues and participants were asked to speak until stopped. As a control task, representing typical cue-based experiments, we analyzed a visually prompted word reading task (Figure 1B; ***Shum et al. (2020)***). While counting and reciting the days or months recruit automatic and continuous speech production, the word reading task follows a structured trial format with distinct silent baseline, stimulus (cue), and speech phases.

**Figure 1.**
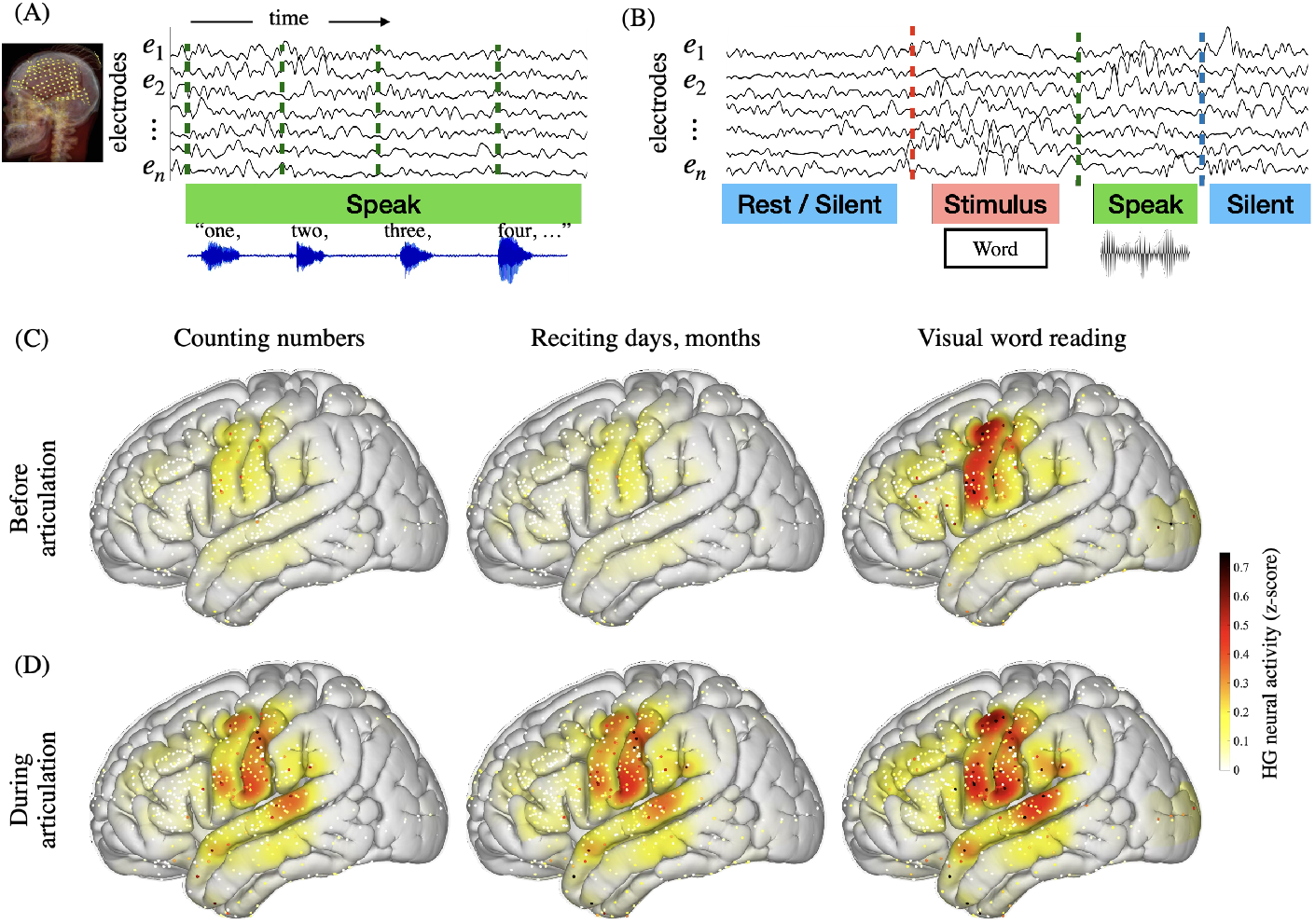
Schematic of speech production tasks and spatial distribution of neural activity. (A) In the automatic speech blocks, participants spoke freely without any cues, prompts, or pauses between words. They were asked to “count from one to twenty,” “recite the days of the week,” and “recite the months of the year.” (B) The cue-based reading task followed a structured trial format. Each trial began with a silent baseline, followed by the visual presentation of a written word. Participants read the word aloud at their own pace, and the next trial began after they finished speaking. High-gamma activity (z-scored) shown averaged across a 500 ms window before (C) and after (D) word articulation onset. Recordings from N=7 participants were time-locked to articulation onset and averaged over trials and corresponding time windows for three speech tasks: counting, reciting days/months, and word reading. Average neural activity is visualized for individual electrodes and as a projection on an average MNI brain.

We analyzed neural activity in the high-gamma broadband range (70-150 Hz) time-locked to articulation onset (see Data collection and general pre-processing). We focused on high-gamma activity due to its strong corelation with underlying spiking neural activity as well as fMRI BOLD signals (***Ray et al. (2008)***; ***Lachaux et al. (2012)***; ***Logothetis (2003)***). The results revealed robust recruitment of a widespread language network before and during speech production (Figure 1 C,D). We observed neural activity in perisylvian areas including superior temporal, post-central, precentral, and frontal regions across all tasks (Figure 1D). However, a clear dissociation was observed between the cued visual word reading and the automatic speech tasks before articulation. The reading task elicited greater neural activity in frontal and pre-central cortices in the 500 milliseconds preceding speech onset compared to the automatic speech tasks (Figure 1C).

To statistically test the involvement of electrodes in speech production across different tasks, we quantified their neural encoding of speech. A direct comparison of high-gamma magnitudes can be misleading due to differences in the task structure as well as the lack of per-trial baseline in automatic speech. We opted for an encoding approach as it circumvents this issue by modeling the relationship between continuous speech output and the neural signal (***Crosse et al. (2016)***). Specifically, we used a multivariate Temporal Response Function (mTRF) model with elastic net regularization (Figure 2; see Neural encoding: mTRF model). Using this approach we characterized, for each electrode, its specific encoding strength based on the correlation (Pearson) between the model’s prediction and the actual neural activity (trained and validated using 5-fold cross-validation; see Neural encoding: mTRF model). Furthermore, the temporal filters learned by the model quantify the specific time-lags at which neural activity is coupled to the produced sound features. This model allows us to directly quantify the encoding dynamics of automatic speech for individual electrodes as well as compare encoding strength across tasks.

**Figure 2.**
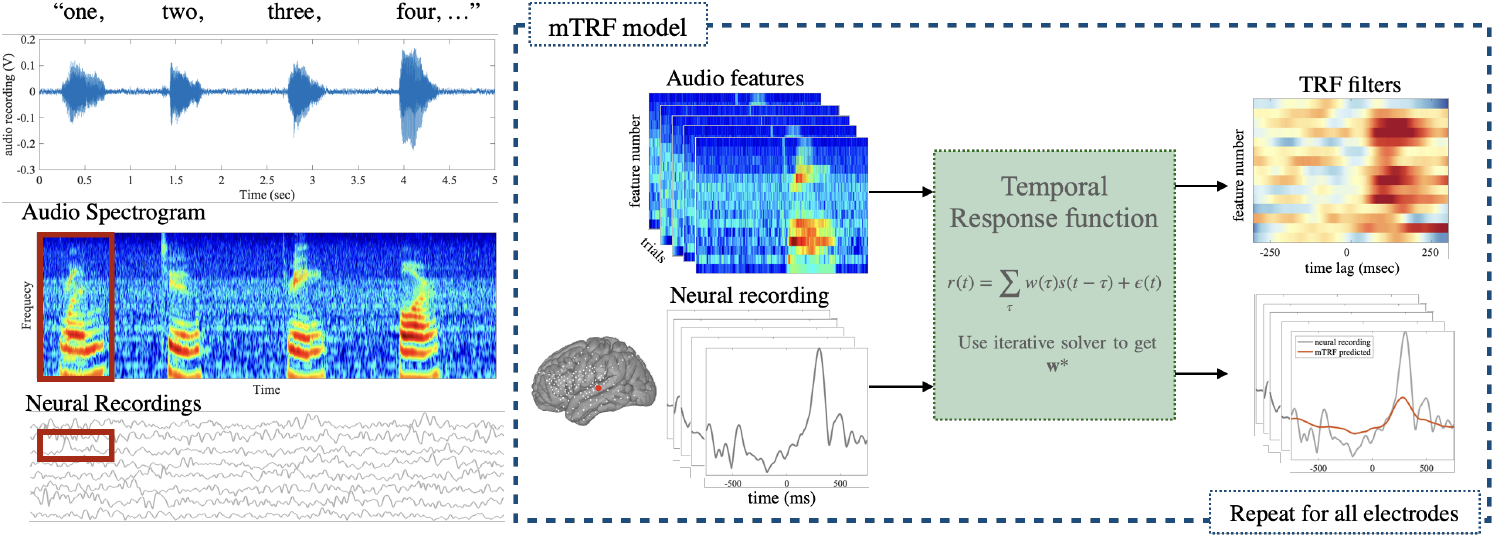
Schematic of the multivariate temporal response functions (mTRF). We use the mTRF model to relate the electrode dynamics to speech audio. We extracted spectrogram features (*s*(*t*)) from the audio waveform. For each electrode, the model predicted the neural responses (*r*(*t*)) with a set of filters (*w*(*τ*)) across various time lags (*τ*). The mTRF filters reveal level of encoding and corresponding timing of neural data relative to speech onset.

The trained models revealed significant speech encoding (permutation test; see Neural encoding: mTRF model) across superior temporal (STG), pre-central (PrG), and post-central gyri for both automatic speech and cue-based reading tasks (Figure 3A). However, the number of electrodes exhibiting significant encoding (see Table 1) as well as the degree of correlation differed markedly in precentral and frontal regions (Figure 3A). To quantify these differences we compared encoding strength across all individual electrodes between tasks (significance assessed using a permutation test, p<0.05 FDR corrected). In contrast to automatic speech, we found significantly higher encoding during word reading, with responses primarily localized across the pre-central and frontal gyri. A minority of electrodes showed higher encoding for automatic speech with post-central gyrus exhibiting a mixture of electrodes encoding speech either for automatic speech or reading task (Figure 3B). This finding provides evidence for neural recruitment across inferior, middle frontal gyri as well as precentral gyrus for cue-based reading task (3B bottom, Wilcoxon signed-rank test *p* = 3.81 × 10^−5^ for frontal and *p* = 2.38 × 10^−7^ for pre-central).

**Table 1.**
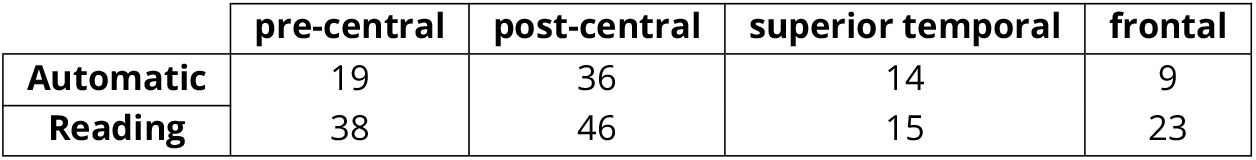
Number of significant encoding electrodes within regions of interest in each task.

|  | pre-central | post-central | superior temporal | frontal |
| --- | --- | --- | --- | --- |
| Automatic | 19 | 36 | 14 | 9 |
| Reading | 38 | 46 | 15 | 23 |

**Figure 3.**
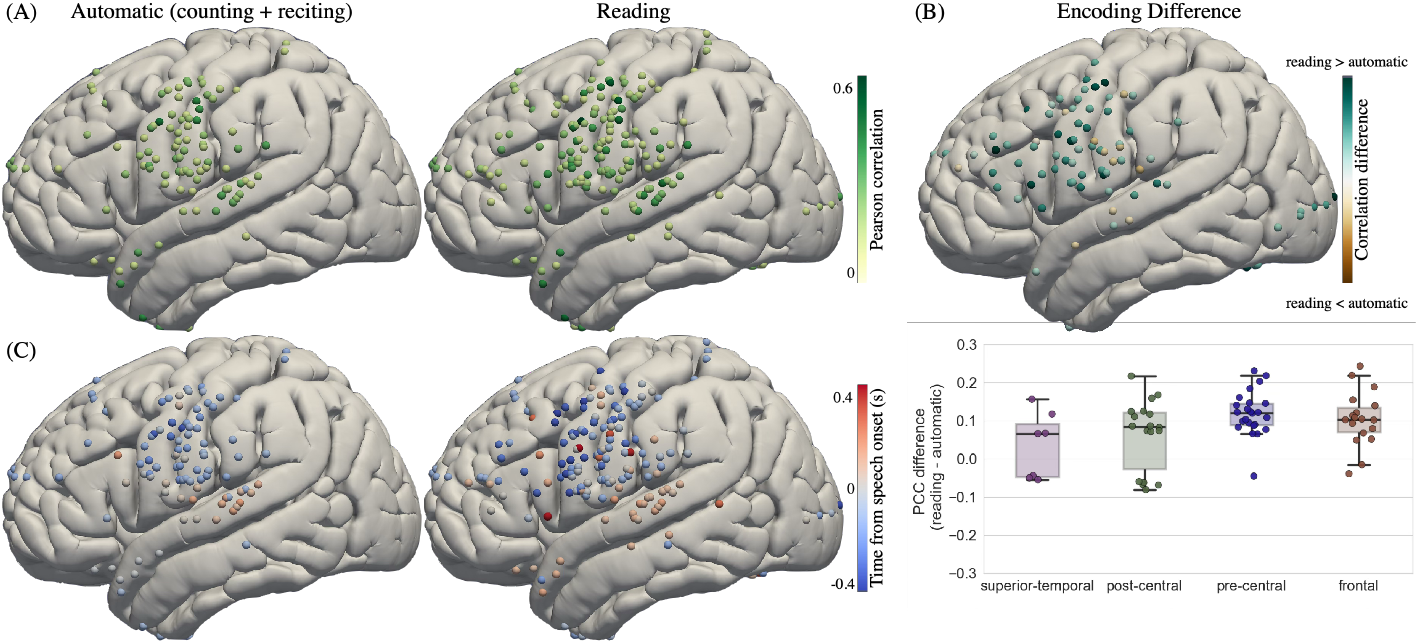
Neural encoding of speech audio features. (A) Encoding strength quantified using the Pearson correlation coefficient for automatic (counting, reciting days and months), and word reading tasks. Electrodes with significant encoding (permutation test, FDR corrected *p* < 0.05) are shown on the MNI brain surface. (B) Comparison of the encoding strength between automatic and cue-based reading task (significance tested against permutation, FDR corrected *p* < 0.05). Top panel: electrodes are color-coded based on encoding correlation difference. Bottom Panel: distribution of correlation difference shown across four regions of interest: pre-central, post-central, middle/caudal superior temporal, and frontal gyri (box plots across electrodes, showing medians and inter-quartile ranges). (C) Peak latencies of the TRFs were computed for electrodes with significant encoding in automatic (counting and reciting days and months), and word reading tasks. Electrodes are color-coded by peak latency relative to articulation onset.

We were interested in characterizing the temporal relationship between speech production and the neural responses across cortices. We derived the peak latency from the encoding model (extremum of the temporal response function) for each significant electrode. During automatic speech, the majority of electrodes in the pre-central and post-central gyri exhibited peak TRF responses before speech onset, whereas electrodes in the STG peaked predominantly after speech onset (Figure 3C). In contrast, during the word reading task, inferior frontal and speech motor cortices showed a more heterogeneous pattern, with TRF peaks occurring both before and after speech onset and spanning a broader range of latencies (Figure 3C).

Many recent studies have focused on decoding speech based on neural data, predominantly acquired during structured or cue-based speech production tasks. We asked if the changes in dynamics seen in our encoding analyses would also be reflected in a brain computer interface decoding approach. We trained a deep neural network (DNN) architecture to predict speech audio from the neural activity (Figure 4 A). Instead of directly decoding to audio waveforms, we use the speech articulatory coding features (SPARC, ***Cho et al. (2024)***) as a latent representation and reconstructed the speech waveform using a pre-trained speech synthesizer. The SPARC model provides an analysis module that maps the original recorded audio to 14 articulatory trajectories (12 related to lip and tongue motion and 2 related to loudness and pitch) and a synthesis module that reconstructs audio from these features. We trained a deep neural network to estimate these articulatory trajectories from the neural signals (see Speech decoding and Figure S1A for details), and during evaluation used the SPARC synthesis module to generate the corresponding audio waveform (Figure 4A). We trained a separate decoding DNN for each participant and evaluated the performance on held-out trials from the counting and reciting days and months tasks. We evaluated model performance via Pearson’s correlation coefficient between the predicted and ground-truth mel-spectrograms. All seven participants showed robust and significant decoding performance relative to a shuffled null model (Wilcoxon rank sum test with FDR correction; Figure 4B; see SI: Decoded audio examples).

**Figure 4.**
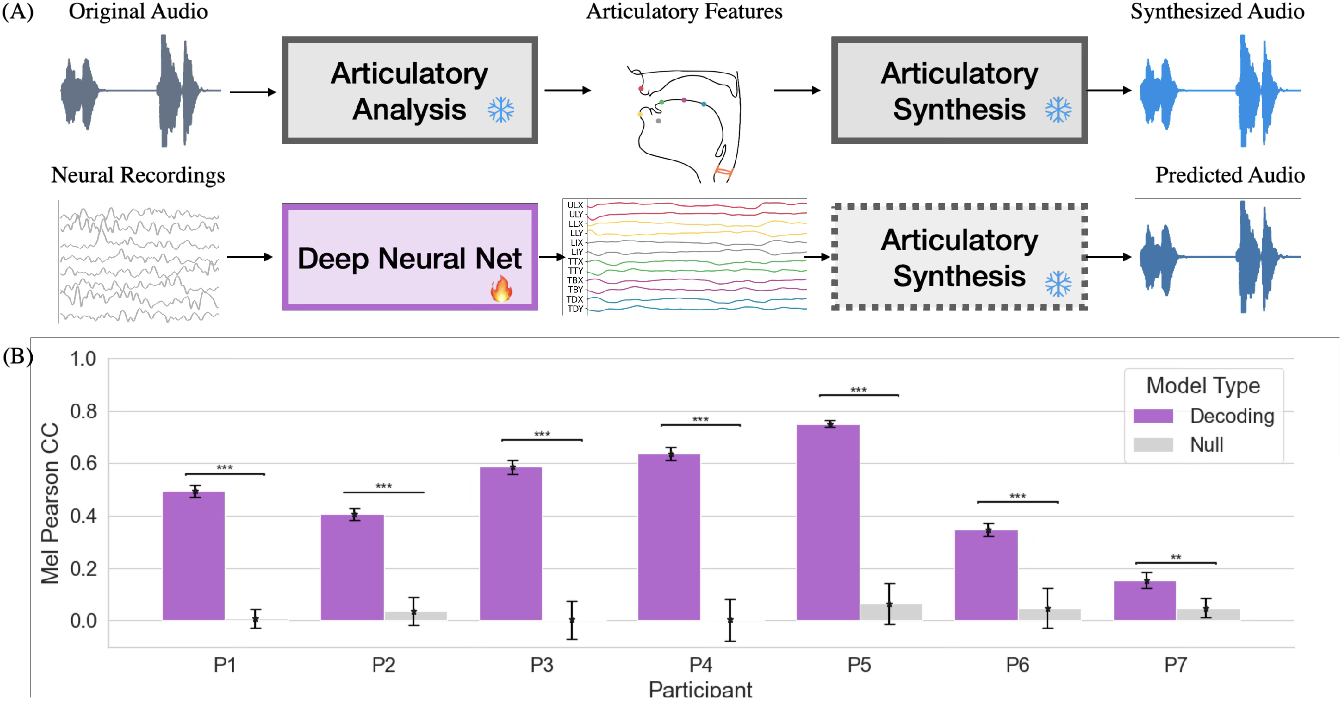
Decoding speech from neural activity via articulatory features. (A) Articulatory trajectories were extracted from the recorded speech using the SPARC model (***Cho et al. (2024)***). The analysis model maps audio waveforms to 14 articulatory features and synthesis model reconstructs audio from these features (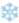shows pre-trained models). A deep neural network (temporal convolution and GRU) was trained to predict articulatory features from neural activity (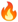shows training from scratch), and the SPARC synthesis module was used to generate predicted audio. (B) Decoding performance quantified by the Pearson correlation (PCC) between predicted and ground-truth mel-spectrograms (bars: mean across held-out trials; error bars: SEM). Significance relative to a shuffled-audio null model was assessed using the Wilcoxon rank-sum test (** : *p* < 0.01, * * * : *p* < 0.001, FDR corrected).

Our encoding analysis provided evidence for mixed latency responses (Figure 3) in speech motor cortex which suggest the presence of both feedforward and feedback representations of speech. To test this, we trained two separate models including strictly feedforward processing (causal) or a common approach in the field that includes both feedforward and feedback processing (non-causal; see Figure S1B for performance comparison). Using an occlusion-based method we were able to quantify the contributions of different electrodes to speech decoding across these models (see Decoding contribution analysis).We measured each electrode’s contribution by the reduction in decoding PCC when the corresponding electrode signal was occluded (that is, we set the signal to zero). We observed that electrodes contributing significantly (p<0.05, Wilcoxon rank sum test) were predominantly localized to the pre-and post-central and superior temporal gyri (Figure 5A). For an identical model trained under a causal setting (only allowing past neural samples to be used), contributing electrodes were predominantly localized to pre- and post-central gyri (Figure 5B). To evaluate the degree to which each electrode is contributing to the non-causal and causal models, we defined a contribution index (ranging from -1 to 1) with positive values indicating higher contribution to non-causal and negative values for the causal model (Figure 5C,D). We observed non-causal contribution (positive index) for electrodes in superior temporal gyrus; as the neural activity related to auditory feedback proceeds articulation. In speech motor areas (pre- and post-central gyri) we observed a mix of both non-causal and causal contributions. Our decoding contributions provide evidence for a mixed feedforward and feedback representation of speech-related activity within both pre-central and post-central gyri in contrast to a purely feedforward control in motor cortex.

**Figure 5.**
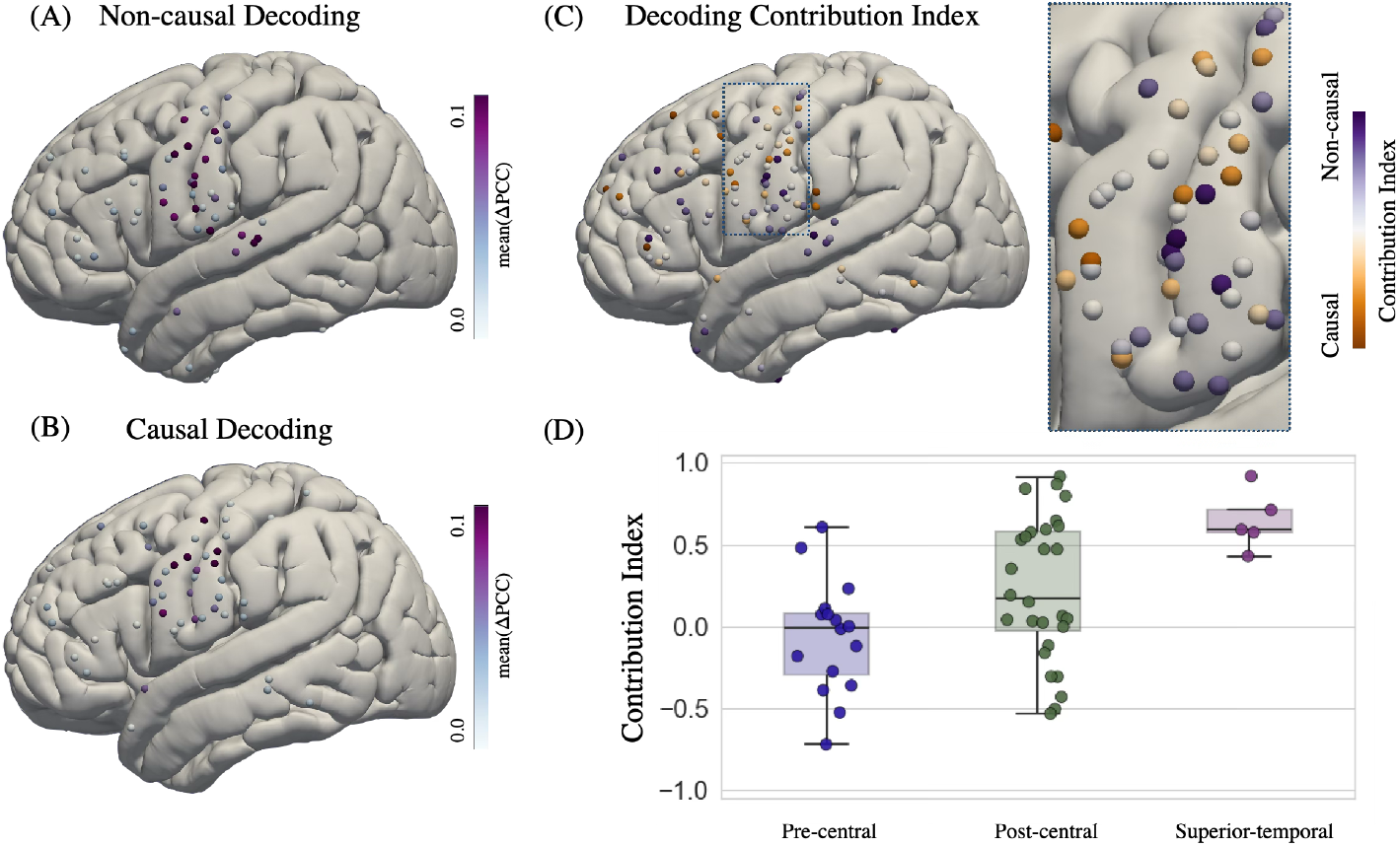
Contribution to speech decoding. Electrodes with significant decoding contributions (color-coded by change in PCC) are displayed for (A) non-causal and (B) causal versions of the DNN speech decoder. (C) The contribution index defined as 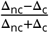 quantifies the level of contribution to the non-causal (positive index) and causal (negative index). (D) The distribution of contribution indices per region-of-interest (box plots show median and quartiles).

## Discussion

We investigated the neural dynamics of automatic speech production using ECoG recordings from the left perisylvian cortex. We examined automatic speech tasks (counting and reciting days or months) alongside a cued word reading task. Our goal was to delineate cortical recruitment during automatic speech production and characterize its temporal dynamics. Using an encoding approach, we quantified the cortical regions engaged during automatic speech and observed significant encoding in ventral speech motor and auditory areas. Relative to the reading localizer, encoding strength was reduced in precentral and inferior frontal gyri. We further demonstrated that automatic speech can be decoded from neural activity using a deep neural network architecture. By constraining the temporal information available to the network during training, we identified electrodes with causal (purely feedforward) and non-causal (mixed feedforward and feedback) contributions to decoding. Taken together, these results delineate the spatiotemporal cortical organization of automatic speech and demonstrate that speech motor cortex supports dynamics beyond purely feedforward control.

The cortical recruitment we observed during automatic speech production is broadly consistent with existing models of speech and prior neuroimaging work (***Birn et al. (2010)***; ***Bookheimer et al. (2000)***; ***Nieder and Mooney (2020)***). The engagement of Rolandic cortex aligns with established accounts of speech motor control and with previous ECoG studies implicating ventral sensorimotor regions in articulation. At the same time, we replicate prior findings that automatic speech is associated with reduced frontal involvement compared to more controlled language tasks (***Bookheimer et al. (2000)***). This pattern has been interpreted as reflecting diminished demands on lexical access and controlled planning during automatic speech relative to cued production (***Gan et al. (2013)***). In line with this interpretation, studies comparing automatic sequences such as counting with controlled verbal fluency tasks report greater involvement of frontal and temporal language regions during controlled speech (***Birn et al. (2010)***). Our ECoG results extend these findings by demonstrating a similar dissociation at the level of cortical encoding strength, with reduced recruitment of precentral and inferior frontal regions during automatic speech production. Above and beyond the existing literature, we established the timing of this recruitment and showed faster timing dynamics in automatic speech compared to the reading task.

While a growing body of work has focused on decoding speech from ECoG and other intracranial signals, these efforts have largely centered on cued speech production tasks (***Chen et al. (2024)***; ***Wilson et al. (2020)***; ***Chakrabarti et al. (2015)***; ***Luo et al. (2022)***; ***Duraivel et al. (2023)***; ***Wyse-Sookoo et al. (2024)***; ***Herff et al. (2015)***). Recent studies have demonstrated high-accuracy neural speech decoding using deep learning frameworks applied to ECoG, including transformer-based and contrastive learning approaches trained on overt or controlled tasks to reconstruct text or speech parameters from cortical signals (***Chen et al. (2024)***; ***Luo et al. (2022)***). We showed that automatic speech can also be decoded significantly using a novel method that leverages pretrained articulatory feature representations (***Cho et al. (2024)***) and a lightweight deep neural network to map neural activity into that feature space. Consistent with our prior work, we further find that the architecture of the decoding model—with access to past and future information (non-causal) versus past only (causal)—shapes the attribution of importance across regions and electrodes. In our previous papers (***Chen et al. (2024)***) we have shown that non-causal models mostly rely on both speech motor and auditory areas, whereas causal models focused predominantly on speech motor cortex in cued or task-based speech production. Here, we observed a similar pattern in automatic speech, and we further showed a mixture of causal and non-causal contributions within pre- and post-central gyri.

Taken together, our results provide a coherent account of the cortical recruitment and dynamics underlying automatic speech production. The convergence of encoding and decoding analyses indicates that automatic speech is not governed by a purely feedforward motor process, but instead reflects an interplay of feedforward and feedback signals within speech motor areas. Our results clarify how automatic speech is instantiated at the cortical level and have practical implications for stimulation mapping in the clinic and neural decoding approaches that aim to capture naturalistic speech behavior.

## Methods

### Participant information

Seven neurosurgical patients (one female; all with left-hemisphere electrode coverage; mean age: 34.57 years, range: 24–55 years) participated in this study after providing written informed consent, which was reaffirmed orally before the experiment began. Electrocorticography electrodes were implanted based exclusively on clinical considerations. Five participants received standard clinical electrode grids with 10 mm inter-electrode spacing (Ad-Tech Medical Instrument, Racine, WI), while two participants consented to implantation of a research hybrid grid (PMT Corporation, Chanassen, MN). The hybrid grids included 64 additional electrodes interleaved among the clinical contacts, maintaining 10 mm spacing overall but incorporating 5 mm spacing over select regions, allowing for denser cortical sampling. The experimental protocol was reviewed and approved by the NYU Langone Medical Center Committee on Human Research.

### Data collection and general pre-processing

ECoG signals were recorded continuously as participants performed the experimental tasks. As an initial quality control step, electrodes exhibiting epileptiform activity, line noise, poor contact quality, or large amplitude drifts were excluded. These exclusion criteria were defined in consultation with the clinical team to remove electrodes that were visually identified as artifacts or outliers. Electrodes containing epileptiform activity were excluded based on clinical characterization as either interictal populations or part of the seizure onset zone. All recordings were re-referenced using the common average reference (CAR) method, in which the mean signal across all electrodes was subtracted from each individual electrode trace. The raw voltage data were then filtered to isolate the high-gamma broadband range (70–150 Hz), and the analytic amplitude (envelope) was computed using the Hilbert transform. The continuous recordings were then segmented into epochs aligned to the onset of speech (articulation-locked).

### Neural activity visualization

When plotting the neural activity (mainly shown in Figure 1), we presented the average z-score. We performed the general pre-processing steps introduced in the previous section. We then normalized each electrode by the mean activity across the task. In Figure 1A-C, the normalized signal for each electrode is averaged over trials and time in a 500 msec window before articulation onset and projected onto a normal MNI brain with a Gaussian kernel of size 50 mm (we restrict the spread of the Gaussian kernel to within the boundaries of the associated cortical region for each electrode; ***Khalilian-Gourtani et al. (2025)***). Similarly, Figure 1D-E shows the average signal in a 500 msec window after articulation onset.

### Neural encoding: mTRF model

We used a multivariate temporal response function (mTRF) implemented with elastic-net regularization to quantify how well the auditory amplitude envelope (stimulus) predicted intracranial electrophysiological signals (responses) on each electrode.

#### mTRF model and fitting procedure

The mTRF model describes how a neural system encodes information related to a given stimulus (***Crosse et al. (2016)***). We model the relation between stimulus and neural response via a linear convolution. Let *r*(*t, n*) denote the neural response sampled at time *t* = 1, *…*, *T*, and recording channel *n* = 1, *…*, *N* and *s*(*t*) be the stimulus. We want to find the channel specific temporal response function *w*(*t, n*), such that

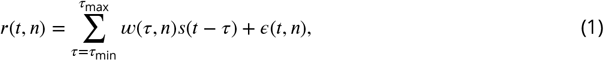

where *τ*_min_ and *τ*_min_ are minimum and maximum filter delays respectively and ε(*t, n*) is the residual not explained by the model. Equivalently, we can form the convolution matrix **S** from the stimulus signal and write (1) in matrix form as

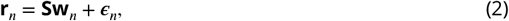

where **r***_n_* = [*r*(1, *n*), *…*, *r*(*T*, *n*)]^⊤^ and other vectors defined similarly (***Crosse et al. (2016)***).

We solve for the TRF, **w**, by minimizing the squared error between the neural response and the prediction from the model. We use elastic net regularization which combines the principles of Ridge (*l*_2_) and Lasso (*l*_1_; which penalizes the absolute value of the coefficients) to enforce both small and sparse TRFs. The mTRF estimation problem with elastic net regularization is written as

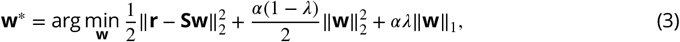

where *α* is the elastic net regularization parameter, and *λ* ∈ (0, 1) controls the ratio of penalizing the *l*_2_ and *l*_1_ terms. We choose *α* = 1, *λ* = 10^−4^, *τ*_min_ = −0.4, and *τ*_max_ = 0.4.

#### Prediction metric and significance against permutation

After fitting, the model predicts the neural responses for held-out test data. Prediction quality for each electrode was quantified as the Pearson correlation coefficient between the predicted and observed time series. To assign statistical significance to observed encoding strengths we built an empirical null distribution of correlation values using a time permutation procedure that preserves the autocorrelation structure of each trial while disrupting the true stimulus–response alignment (***Flinker et al. (2019)***). For each permutation step, we generated a permuted response dataset by applying an independent random circular shift to each trial’s response time series:

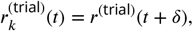

where *δ* is drawn uniformly from {0, …, *T* − 1}. This operation preserves within-trial temporal structure but destroys the precise stimulus to response temporal alignment. We then fitted a model for each permutation and computed predictions and correlation values using the same training and testing pipeline, yielding a permuted correlation for all electrodes that had valid observed correlations. We then aggregated the permuted correlations to form a null distribution at each electrode. From this null distribution we compute the significance for each electrode. Finally, to correct for multiple comparisons across electrodes, the resulting *p*-values were adjusted using Benjamini/Hochberg false discovery rate (FDR) at *α* = 0.05. To test whether encoding level in an electrode was significantly different between tasks, we compared the difference in Fisher-z transformed correlation coefficients between tasks against a permutation derived null distribution and assessed significance with two-sided tests followed by FDR correction across electrodes.

#### Temporal-latency characterization of filters

To characterize the latency of the TRF for each significant electrode we averaged the fitted weights across filters (for multiple stimulus features) to produce a single lag profile per electrode. Local extrema in the lag profile were identified. If a single local extrema was found it was taken as the TRF peak latency, if multiple peaks were present the extrema with the largest absolute weight was chosen.

### Speech decoding

#### Speech articulatory coding

We used the pretrained Speech Articulatory Coding (SPARC) model (***Cho et al. (2024)***) to extract articulatory features from the recorded speech sounds. We chose SPARC because its articulatory representation provides a compact and interpretable feature space, which we could map to it from the neural activity and use the pretrained synthesizer to generate decoded speech. For each participant and each spoken word in the counting and recitation (days and months) tasks, we extracted the articulatory trajectories using the pretrained SPARC analysis (coder) model. Before feature extraction, we normalized the audio amplitude and resampled it to 16 kHz. Features were computed from 280 ms before word onset to at least 1000 ms after onset; in rare case where the word’s duration exceeded 1000 ms, a longer window was used. The SPARC representation consists of 12 temporal signals capturing articulator movements (e.g., lip and tongue positions) and two additional temporal signals representing pitch and loudness. All articulatory features generated by SPARC were sampled at 50 Hz.

#### Deep neural network decoding architecture

We extracted the neural activity corresponding to each audio trial by computing the analytic amplitude of the high-gamma (70-150 Hz) broadband signal (with similar pre-processing steps as in Data collection and general pre-processing). The neural data were then downsampled from 512 Hz to 125 Hz, following our prior speech decoding work (***Chen et al. (2024)***; ***Wang et al. (2023)***). We designed a deep neural network to map from the neural activity to the articulatory features. The model receives a window of neural signals *r*(*t, n*) and predicts the articulatory features *h*(*t, m*), where *m* indexes the 14 feature dimensions. Because the sampling rates of the neural and articulatory domains differ, we first applied linear temporal convolution layers (transpose conv to upsample and conv to downsample) and extracted resampled temporal features matching the 50 Hz rate of the articulatory features. The temporally aligned features were then passed through a gated recurrent unit (GRU), which transformed the sequence into a latent representation. Finally, we used a multilayer perceptron (MLP) to predict the articulatory features from the GRU outputs at each timestep. The model architecture details are presented in Table 2. To make a causal version of this architecture, we switched the convolution modules for causal convolution (***Chen et al. (2024)***) and used a uni-directional GRU instead of a bi-directional one.

**Table 2.** Network architecture and parameter settings for the neural-to-articulatory decoder.

| Component | Description / Parameters |
| --- | --- |
| Input neural signal | high-gamma analytic amplitude, downsampled to 125 Hz. |
| Temporal convolution | 1D linear transpose convolution, kernel size $k = 44$; stride $s = 2$. |
| Temporal convolution | 1D linear convolution, kernel size $k = 1$; stride $s = 5$. |
| GRU layer | 3-layer GRU applied to temporally aligned features; hidden size $d = 512$. |
| MLP decoder | Fully connected MLP; input $d = 512$, hidden $d = 256$, output $d = 14$; ReLU nonlinearity |
| Training loss | Mean squared error (MSE) and Huber loss between predicted and ground-truth articulatory features |
| Optimization | Adam optimizer, learning rate $3 \times 10^{-4}$, batch size 64. |
| Causality | non-causal model uses both past and future context (non-causal conv and bi-directional GRU); Causal model uses only past neural context (causal conv and uni-directional GRU) |

#### Decoding contribution analysis

For each participant and each trained decoding model (non-causal or causal), we first computed the Pearson correlation coefficient between the ground-truth articulatory features and the model’s predictions on the held-out test set. For each electrode, we then zeroed out its signal in the test data, recomputed the predicted features, and calculated the resulting PCC. We tested whether the PCC after occlusion differed significantly from the full model (Wilcoxon rank-sum test with FDR correction). For electrodes that showed a significant effect, we reported the mean difference in PCC between the full and occluded models. We defined a contribution index defined as 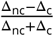 to quantify the level of contribution to the non-causal (Δ_nc_, positive index) and causal (Δ_c_, negative index) models.

## Acknowledgments

This work was supported by National Institutes of Health grants R01NS109367, R01NS115929, and R01DC018805 (A.F.) and National Science Foundation IIS-2309057 (A.F., Y.W.).

## Code availability

The code for processing and analysis of the encoding and decoding are provided publicly here: https://github.com/amirhkhalilian/automatic_speech. A standalone installable version of the encoding model is provided here: https://github.com/amirhkhalilian/ADMM_mTRF.

## Data availability

The dataset analyzed for this study is available from the corresponding authors upon reasonable request.

## Supplemental Information

### SI: Speech decoding

**Figure S1.**
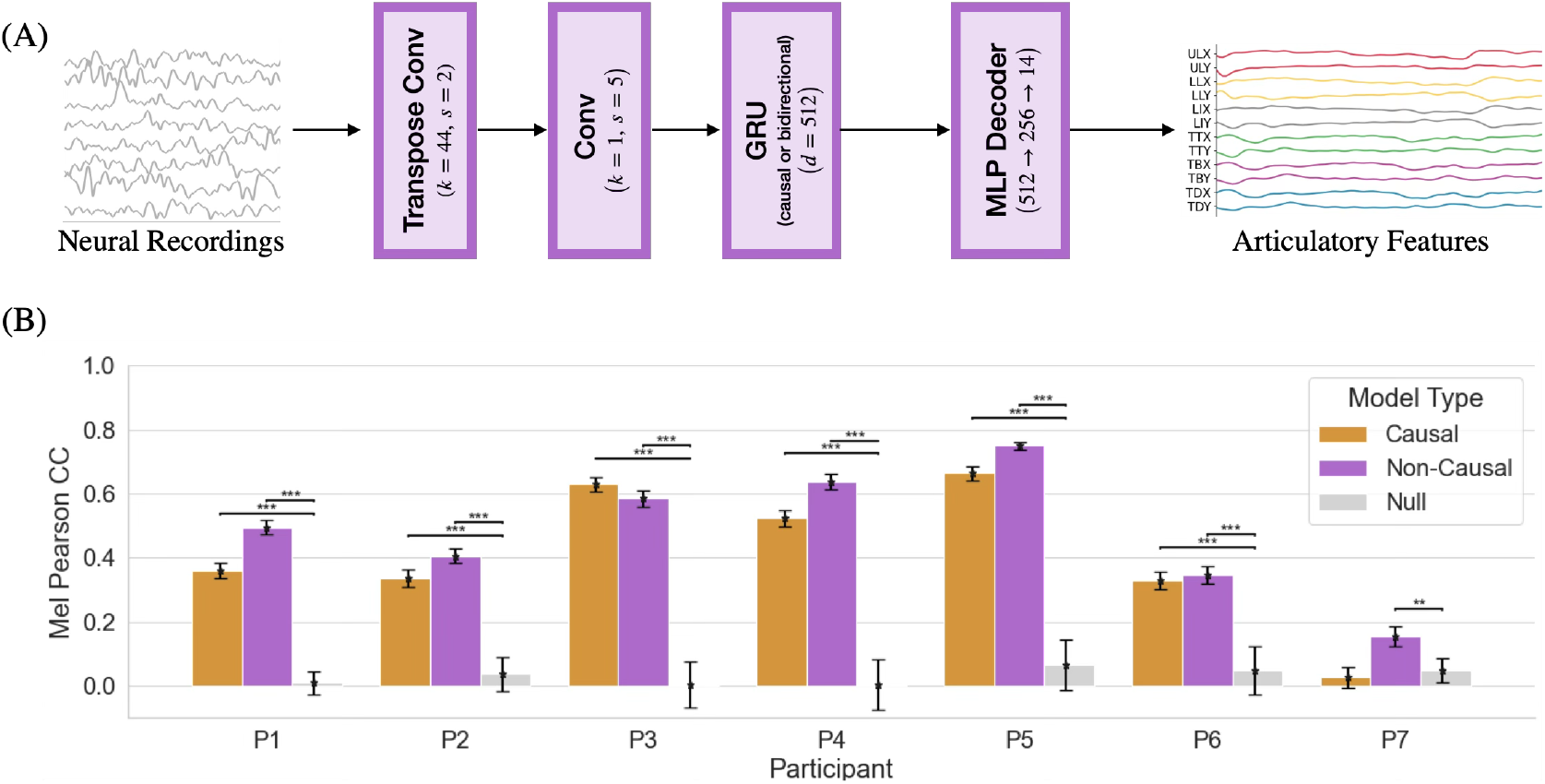
Speech decoding neural network. (A) The architecture of our deep neural network (temporal convolution and GRU) designed to predict articulatory SPARC features from neural activity. In the causal setting, we used causal convolution and unidirectional causal GRU. (B) Decoding performance quantified by the Pearson correlation (PCC) between predicted and ground-truth mel-spectrograms for Non-causal and causal decoding architectures (bars: mean across held-out trials; error bars: SEM). Significance relative to a shuffled-audio null model was assessed using the Wilcoxon rank-sum test (** : *p* < 0.01, * * * : *p* < 0.001, FDR corrected).

### SI: Decoded audio examples

Audio decoding examples are provided in the attached HTML file (`audio_player.html`). For each example, we include the original audio recording, along with the decoded audio waveforms generated by the deep neural network using both non-causal and causal architectures. Decoded speech is synthesized from the predicted SPARC feature representation using the provided synthesizing vocoder that conditions on speaker identity. Because the original speaker identity is not directly available to the vocoder (i.e. outside its original training set), audio is generated using the closest matching voice. In addition, we include an example of a failure case in which part of the decoded audio is incorrectly mapped to a different phoneme, resulting in speech that falls outside the target vocabulary.

