## Supplementary material for "Neural Dynamics of Automatic Speech Production": audio_player.html

Automatic Speech Decoding

### Speech Decoding Pipeline

Many recent studies have focused on decoding speech based on neural data, predominantly acquired during a structured cue-based task. We asked if automatic and overlearned speech, such as counting or reciting name of days, can also be decoded from neural activity. We trained a novel deep neural network (DNN) architecture to predict speech audio from the neural activity. Our DNN is trained to estimate the articulatory trajectories of SPARC from the neural signals, and during evaluation
we used the SPARC synthesis module to generate the corresponding audio waveform. The following examples show the original audio and the decoded waveforms using non-causal and causal DNN architectures.

### Decoding Audio Examples

Original Audio

Non-causal Decoding

Causal Decoding

Original Audio

Non-causal Decoding

Causal Decoding

Original Audio

Non-causal Decoding

Causal Decoding

### Failure Case Example

Original Audio

Non-causal Decoding

Causal Decoding
