## Supplementary material for "Neural Dynamics of Automatic Speech Production": SI_Automatic_Speech_Khalilian_2026.pdf

### Supplemental Information

#### SI: Speech decoding

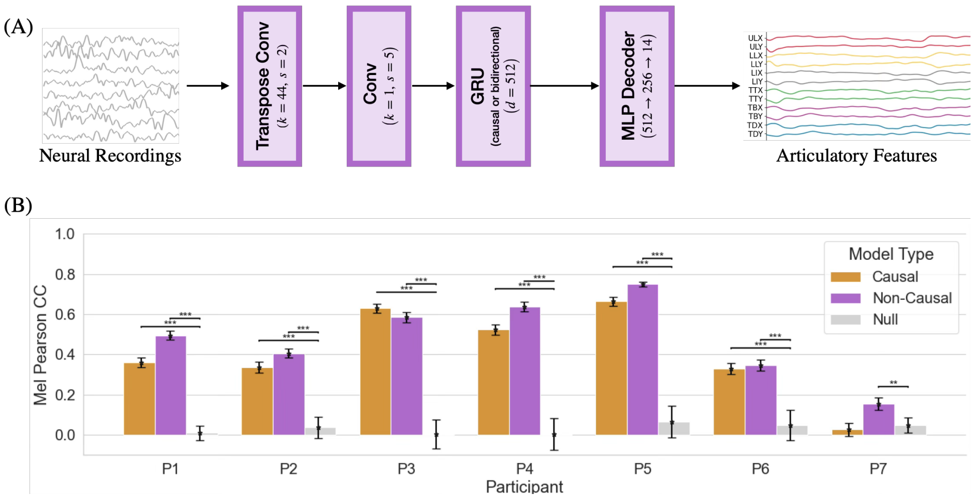


**Figure S1. Speech decoding neural network**

(A) The architecture of our deep neural network (temporal convolution and GRU) designed to predict articulatory SPARC features from neural activity. In the causal setting, we used causal convolution and unidirectional causal GRU. (B) Decoding performance quantified by the Pearson correlation (PCC) between predicted and ground-truth mel-spectrograms for Non-causal and causal decoding architectures (bars: mean across held-out trials; error bars: SEM). Significance relative to a shuffled-audio null model was assessed using the Wilcoxon rank-sum test ($** : p < 0.01$, $*** : p < 0.001$, FDR corrected).

#### SI: Decoded audio examples

Audio decoding examples are provided in the attached HTML file (`audio_player.html`). For each example, we include the original audio recording, along with the decoded audio waveforms generated by the deep neural network using both non-causal and causal architectures. Decoded speech is synthesized from the predicted SPARC feature representation using the provided synthesizing vocoder that conditions on speaker identity. Because the original speaker identity is not directly available to the vocoder (i.e. outside its original training set), audio is generated using the closest matching voice. In addition, we include an example of a failure case in which part of the decoded audio is incorrectly mapped to a different phoneme, resulting in speech that falls outside the target vocabulary.
